# MaxQuant.Live enables global targeting of more than 25,000 peptides

**DOI:** 10.1101/443838

**Authors:** Christoph Wichmann, Florian Meier, Sebastian Virreira Winter, Andreas-David Brunner, Jürgen Cox, Matthias Mann

## Abstract

Mass spectrometry (MS)-based proteomics is often performed in a shotgun format, in which as many peptide precursors as possible are selected from full or MS1 scans so that their fragment spectra can be recorded in MS2 scans. While achieving great proteome depths, shotgun proteomics cannot guarantee that each precursor will be fragmented in each run. In contrast, targeted proteomics aims to reproducibly and sensitively record a restricted number of precursor/fragment combinations in each run, based on pre-scheduled mass-to-charge and retention time windows. Here we set out to merge these two concepts by a global targeting approach in which an arbitrary number of precursors of interest are detected in real-time, followed by standard fragmentation or advanced peptide-specific analyses. We made use of a fast application programming interface to a quadrupole Orbitrap instrument and real-time recalibration in mass, retention time and intensity dimensions to predict precursor identity. MaxQuant.Live is freely available (www.maxquant.live) and has a graphical user interface to specify many pre-defined data acquisition strategies. Acquisition speed is as fast as with the vendor software and the power of our approach is demonstrated with the acquisition of breakdown curves for hundreds of precursors of interest. We also uncover precursors that are not even visible in MS1 scans, using elution time prediction based on the auto-adjusted retention time alone. Finally, we successfully recognized and targeted more than 25,000 peptides in single LC-MS runs. Global targeting combines the advantages of two classical approaches in MS-based proteomics, while greatly expanding the analytical toolbox.

## INTRODUCTION

Mass spectrometry (MS)-based proteomics has matured into a versatile and widely used analytical tool in the life sciences[1]–[3]. State-of-the-art workflows cover the proteome of model organisms to near-completeness and sensitivity extends into the attomole range[4]–[7]. Still, many, and in particular low-abundance, proteins escape accurate and reproducible quantification across large sets of biological samples, which hinders wider applications of proteomics in systems biology and translational medicine[8], [9]. In part, this is due to the complexity of bottom-up proteomics samples. Enzymatic digestions of protein extracts are comprised of millions of peptide species and even with liquid chromatography, many of them co-elute, while spanning several orders of magnitude in abundance[10]. In data-dependent acquisition schemes (DDA), the mass spectrometer acquires as many MS2 spectra as possible to maximize the number of peptide identifications. As high-abundance ions are more likely to yield high-quality MS2 spectra, precursors are typically prioritized for isolation and fragmentation by their abundance and dynamic exclusion is employed to prevent their re-sequencing. This topN approach has been the method of choice for unbiased and comprehensive proteomic studies for many years. However, given the enormous number of precursor candidates and unavoidable run-to-run variabilities, a specific peptide may not be fragmented in every run. Data-independent acquisition (DIA) aims to avoid stochasticity by repeatedly cycling through fixed isolation windows[11], however, it increases spectral complexity and may diminish dynamic range, and is not subject of this manuscript.

In contrast to DDA and DIA, the goals of targeted proteomics methods are to analyze a limited number of selected proteins of interest with maximum sensitivity and reproducibility[12]. Rather than selecting precursors based on the MS1 scans, the instrument is instructed to continuously fragment certain pre-defined precursor-fragment mass combinations during scheduled time windows, in which the targeted peptides are expected.

Traditionally, these experiments have been performed on triple quadrupole instruments, even before the advent of proteomics. However, today high-resolution time-of-flight or Orbitrap mass analyzers are gaining popularity. Instead of recording one or a few specified precursor-fragment transition, these instruments acquire complete MS2 spectra and therefore monitor all fragment ions simultaneously, which increases specificity and quantitative accuracy[13], [14]. In practice, setting up a robust targeted proteomics experiment with a desired coverage remains challenging as the number of targets needs to be balanced with acquisition speed and sensitivity[15], [16]. Selecting too many targets may unduly reduce the acquisition time for each of them, while specifying narrow LC elution windows increases the risk of missing a peptide entirely as the retention times cannot be estimated very accurately beforehand. Reports in the literature generally employ minute-wide monitoring windows to target tens and sometimes hundreds peptides and proteins. Furthermore, despite the creation of community-wide MRM peptide libraries[17]– [19], these assays are typically reestablished and optimized in each laboratory.

To address some of the above limitations, Domon and co-workers spiked-in isotope-labeled variants of the peptides of interest to trigger ‘pre-targeting’ and targeting events more precisely[20]. Coon and colleagues used the expected elution order of peptides to bias DDA toward peptides of interest[21]. Building on the MaxQuant software suite[22], [23] our own group developed the MaxQuant-RealTime framework, which identified peptides within milliseconds, providing a basis to implement intelligent data acquisition methods in different research scenarios[24]. Although these concepts and potential applications are very promising in principle, the uptake of the underlying software packages was limited.

Proteomics post-processing algorithms generally contain a mass re-calibration step as well as retention time alignment, which can be used to transfer identifications between runs[22], [25], [26]. For instance, MaxQuant can achieve sub-parts-per-million (ppm) mass accuracies and absolute retention time deviations below 30s by non-linear re-calibration and alignment. We reasoned that approaching such an accuracy in real-time could dramatically improve our ability to predict the appearance of a very large number of peptides. This would drastically reduce the monitoring time for each peptide, and might allow extending the targeting concept to a global, proteome-wide scale.

To realize this vision, we developed the freely available software MaxQuant.Live, which interacts with any Thermo Fisher Q Exactive mass spectrometer via the re-designed instrument application programing interface (IAPI). Scan modules (‘apps’) can be plugged into the MaxQuant.Live core application on the acquisition computer, allowing straightforward implementation and modification of standard acquisition schemes as well as advanced data acquisition strategies based on live data analysis.

## EXPERIMENTAL PROCEDURES

### Cell culture and sample preparation

We cultured the human HeLa cancer cell line (S3, ATCC) in Dulbecco’s modified Eagle’s medium (DMEM) with 10% fetal bovine serum, 20 mM glutamine and 1% penicillin-streptomycin added (PAA Laboratories). Metabolic stable isotope labeling was performed in arginine- and lysine-free DMEM, fortified with arginine and lysine with natural isotope abundances (light channel) or stable-isotope labeled arginine-10 and lysine-8 (Cambridge Isotope Laboratories) as previously described[27]. Cells were collected by centrifugation, washed twice with cold phosphate-buffered saline, pelleted and stored at −80 °C.

We lysed the cells, and reduced and alkylated the proteins in a single reaction vial with sodium deoxycholate (SDC) buffer containing chloroacetamide (PreOmics GmbH) similar to our previously published protocol[28]. Cells were suspended in the SDC buffer and boiled for 10 min at 95 °C. To disrupt remaining cellular structures and shear nucleic acids, we sonicated the suspension for 15 min at full power (Bioruptor, Diagenode). The crude protein extracts were enzymatically digested with LysC and trypsin (1:100, enzyme wt: protein wt) overnight at 37 °C before stopping the reaction with 5 volumes of isopropanol/1% trifluoroacetic acid (TFA). Peptide micro-purification and de-salting was performed on styrenedi-vinylbenzene-reversed phase sulfonate StageTips. Following sequential washing steps with isopropanol/1% TFA and water with 0.1% TFA, peptides were eluted with 80% acetonitrile (ACN) containing 1% ammonia. The vacuum dried eluates were reconstituted in water with 2% ACN and 0.1% TFA for further analysis.

### Liquid Chromatography and Mass Spectrometry (LC-MS)

In single LC-MS runs, approximately 500 ng of purified whole-cell digests were analyzed with an EASY-nLC 1200 nanoflow chromatography system (Thermo Fisher Scientific) coupled online to a hybrid quadrupole Orbitrap mass spectrometer (Thermo Q Exactive HF-X[29]). The peptides were separated at 60 °C on a 50 cm long column (75 μm inner diameter) packed with 1.9 μm porous silica beads (Dr. Maisch GmbH), and electrosprayed from a laser-pulled silica emitter tip at 2.4 kV. Mobile phases A and B were water with 0.1% formic acid (vol/vol) and 80/20/0.1% ACN/water/formic acid (vol/vol/vol). To elute the peptides at a constant flow rate of 300 nL/min, a binary gradient was ramped from 5% to 30% B within 95 min, followed by an increase to 60% B within 5 min and further to 95% B for washing. After 5 min, the organic content was decreased to the starting value within 5 min and the column was re-equilibrated for another 5 min.

Standard top15 DDA methods were generated with the graphical Thermo Xcalibur method editor. Full MS scans in the mass range from *m/z* 300 to 1650 were acquired with a 128 ms transient time corresponding to a resolution of 60,000 at *m/z* 200. The target value for the automatic gain control (AGC) algorithm was set to 3 x 10^6^ charges, which was typically reached within about 1 ms while peptides were eluting. Precursor ions for MS2 scans were isolated with a +-0.7 Th window centered on the precursor mass and fragmented with higher energy collisional dissociation (HCD)[30] at a normalized collision energy of 27. MS2 spectra were acquired with a resolution of 15,000 at *m/z* 200, and the maximum ion injection time and the AGC target were set to 25 ms and 1 x 10^5^ charges, respectively. Only precursors with assigned charge states >= 2 and <= 5 were considered, and previously sequenced precursors were dynamically excluded for 30 s.

### Acquisition software

MaxQuant.Live (Version 0.1) was continuously running in its ‘listening mode’ on the acquisition computer waiting for a signal to load and execute a scan protocol from the library. For scheduling LC-MS run, we used the sequence list from Xcalibur whose entries contain settings for the LC device as well as the method for the mass spectrometer. While using the normal LC-settings we constructed the instrument method in such a way that it encodes the start signal for MaxQuant.Live to load a given scan protocol and take over the control of the mass spectrometer for the whole run upon starting.

Scan protocols for the different targeting strategies were all specified using the targeting app that is included in MaxQuant.Live. As initial settings for peptide recognition, we chose by default a mass tolerance of ±10 ppm, a retention time tolerance of ±3 min and an intensity threshold value of 10^-5^ from the expected intensity of the target. To calculate the corrections, the adaptive correction includes the peptides recognized within the last 3 minutes but retains a minimum of the last 100. The correction automatically started 6 min after the first peptide was recognized and the mass tolerances were set to 6.0 ppm and peptide elution tolerances to the triple of the standard deviation.

### Breakdown curves

As an example of advanced acquisition schemes enabled by MaxQuant.Live, we studied the large-scale and automated acquisition of HCD fragmentation characteristics of peptides. In triplicate runs of a HeLa digest, we targeted 1,000 precursors, using 10,000 endogenous peptides for real-time corrections. Upon recognition in real-time, precursors were isolated with a +-0.2 Th window and repeatedly fragmented with increasing collision energy in ten steps from the normalized collision energy (NCE) 16 to 34. Other than that, the MS parameters were set as above. Target peptides were selected randomly from the top 50% abundance quantile of peptides identified in a standard DDA run of the same digest after removal of contaminants and reverse hits and filtering for the most abundant evidence of each unique peptide sequence. The tolerances for the real-time correction were the default values listed before.

### Predictive Multiplexed Selective Ion Monitoring (pmSIM)

Light and heavy labelled tryptic Hela lysates were mixed in a ratio of 4:1 and 500 ng were injected on column. DDA raw files were analyzed using MaxQuant to identify light to heavy SILAC[31] peptide ratios. To generate a targeting list for MaxQuant.Live, the ‘evidence’ output file was filtered for modified sequence duplicates, missed cleavages, keeping only unmodified peptides with a sequence length less than 25 amino acids and a retention length less than 2 min. Peptides for retention correction were additionally filtered for >10 and < 90 min retention time, after which the top 10,000 most intense light channel peptides were selected. The fifty peptides for selected ion monitoring (SIM) were randomly chosen from a list fulfilling the following criteria: retention time 20-80 min, no reported L/H ratio, an intensity of zero in the heavy channel. The initial retention time tolerance was ±10 min and the final value 1.5 fold of the elution time standard deviation. The tolerances for the real-time correction were set to the default values listed above.

MaxQuant.Live pmSIM experiments were performed with a 1 Th isolation window and a +0.2 Th offset and acquired with a resolution of 120,000 at *m/z* 200. The heavy and light channels were multiplexed in a single scan. A maximum of 1 x 10^5^ ions were collected in each channel with a maximum ion injection time of 48 and 192 ms for the light and heavy channel, respectively.

Data analysis of the pmSIM experiment was performed with the Skyline[32] (Version 4.1.0.18169) and XCalibur (3.1.66.10) software suites. The SIM targeting raw file was split into SIM and MS1 scans and analyzed independently.

### Global targeting

To build a reference DDA dataset, 500 ng of tryptic Hela digest were measured in triplicate and raw files were analyzed with MaxQuant. The matching between runs feature was activated using the default settings. Peptide identities as well as their mass, charge state, retention time and intensity were extracted from the evidence output file and used to generate targeting lists for MaxQuant.Live. Only peptides with a retention time between 10 and 100 minutes that were identified by MS/MS or matching in all three replicates were candidates for the targeting lists and any hits from the reverse decoy library and potential contaminants were excluded from the selection. To generate the targeting lists, 100, 1,000, 5,000, 10,000, 20,000 or 30,000 peptides were randomly selected from all peptides fulfilling the above criteria, ensuring a uniform distribution of targets over the whole abundance range. For real time correction, we selected the 10,000 most abundant peptides identified by MS/MS or matching in all three replicates with a retention length less than 30 s. The tolerances for real-time correction were the default values listed above. To demonstrate the functionality of the real-time correction we performed an additional run with 20,000 targeting peptides, in which the minimal mass tolerance was 4.5 ppm and the retention time windows size was 2.5 fold of the standard deviation of the peptide elution times. Here, the correction was calculated from all the peptides that were recognized within the last minute but at least the last 20.

Full MS scans in the mass range from *m/z* 300 to 1650 were acquired with a resolution of 60,000 at *m/z* 200 and an automatic gain control (AGC) target of 3 x 10^6^. Target peptides had a ‘Life Time’ (max. time between recognition and fragmentation) of 1000 ms and were isolated for MS2 scans with a +-0.2 Th window centered on the precursor mass and fragmented with at a NCE of 27. MS2 spectra were acquired with a resolution of 15,000 at *m/z* 200, and the maximum ion injection time and the AGC target were 110 ms and 1 x 10^5^ charges, respectively. For real time correction we used the default values mentioned before. Raw files of the targeting runs were analyzed in MaxQuant together with the standard DDA runs with the matching between runs feature activated.

### Proteomics data processing

*S*hotgun proteomics raw data acquired with either the standard user interface or MaxQuant.Live were processed with MaxQuant[22] (version 1.6.1.13) using the default settings if not stated otherwise. The built-in Andromeda search engine[23] scored MS2 spectra against fragment masses of tryptic peptides derived from a human reference proteome containing 95,057 entries including isoforms (UniProt, release 2018/06) and a list of 245 potential contaminants. We required a minimum peptide length of 7 amino acids and limited the search space to a maximum peptide mass of 4,600 Da and two missed cleavage sites. Carbamidomethylation of cysteine was specified as a fixed modification, and methionine oxidation and acetylation at protein N-termini as variable modifications. MaxQuant uses individual mass tolerances for each peptide, while the initial maximum precursor mass tolerances were set to 20 ppm in the first search and 4.5 ppm in the main search, and the fragment mass tolerance was set to 20 ppm. The false discovery rate was controlled with a target-decoy approach at less than 1% for peptide spectrum matches and less than 1% for protein group identifications.

### Bioinformatics

Post-processing was performed in either Perseus[33], the R computational environment[34] or the Python programming language. Potential contaminants, reverse database hits and proteins identified by only one modified peptide were excluded from the analysis.

## RESULTS

Here we describe the development of a software framework termed MaxQuant.Live for real-time monitoring of mass spectrometric data and controlling of the data acquisition. We demonstrate its usability and performance in terms of scanning speed using a re-implemented topN method. We demonstrate that thousands of peptides of interest can be detected and immediately selected for deeper analysis, greatly extending the toolbox for targeted proteomics. To explore the current limits of our technology, we targeted over 20,000 peptides in a single experiment.

### Design and functionality of MaxQuant.Live

A few years ago, Thermo Fisher Scientific developed an IAPI that enables fast, bidirectional communication between a Q-Exactive mass spectrometer and an outside. We developed a software module, written in the C# programming language, containing functionality for advanced data acquisition and analysis in real-time, which communicates with the mass spectrometer through the IAPI (EXPERIMENTAL PROCEDURES). We termed the program MaxQuant.Live because it forms a bridge between intelligent data acquisition and downstream analysis in the MaxQuant environment. In one direction, the IAPI transmits every measured mass spectrum to our software on the fly and in the other direction it enables MaxQuant.Live to send scan commands to the instrument every time it is ready to accept new instructions. **Figure 1** illustrates the interplay between the core module of MaxQuant.Live and the mass spectrometer enabled by the IAPI. The engine in MaxQuant.Live executes a run-specific scan protocol (see below) which contains the acquisition strategy for the current LC-MS run and which is loaded from the scan protocol library. The scan requests generated by the engine are collected in the local scan queue before they are pushed sequentially to the MS instrument.

**Figure 1.**
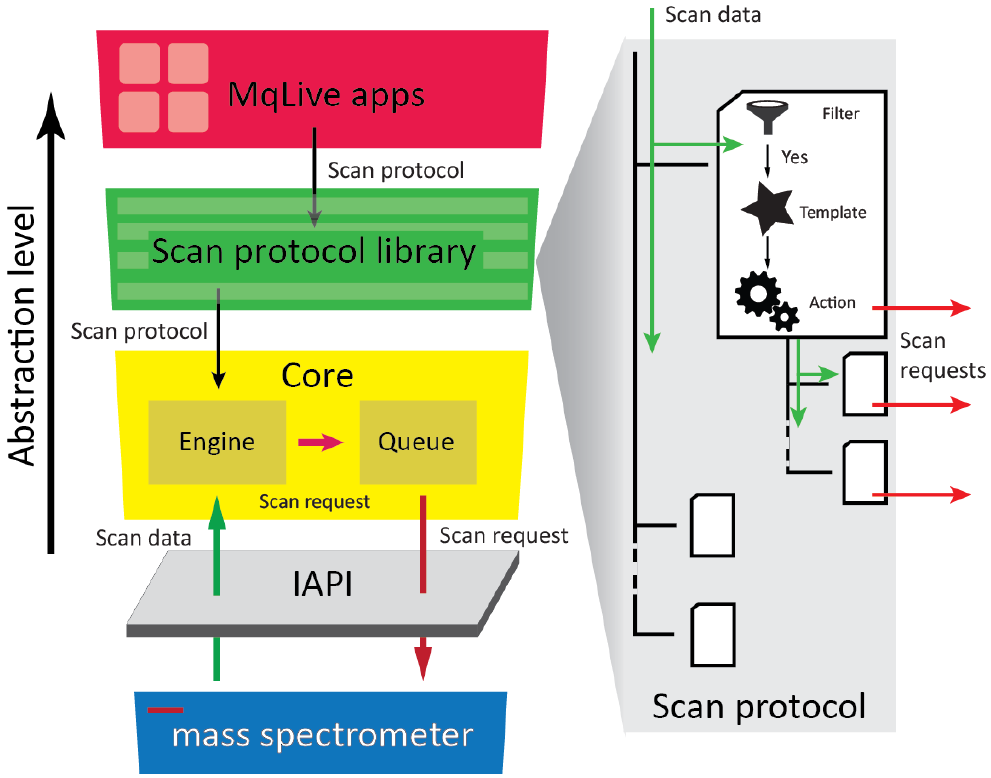
Architecture of MaxQuant.Live and the logic of its scan protocols. The core of our software (yellow box) handles the real-time control of the mass spectrometer using the IAPI by Thermo Fisher. Its engine processes the scan data according to a scan protocol, which specifies a data-acquisition strategy trough a decision tree logic (right). Scan protocols for different applications are collected in a library and can be generated by small applications (‘apps’).

In case the scan queue is empty it periodically sends fallback scan requests to prevent the instrument from running idle or changing its operation status. This design of the scan queue ensures that the core module of MaxQuant.Live keeps control of the instrument during the entire run once a scan protocol has been loaded from the library and while the instrument is connected to the IAPI. Due to this generic additional abstraction layer, our core module is independent of the attached IAPI and could also be combined with instrument control libraries of other mass spectrometers.

The scan protocol specifies the acquisition concept for a particular LC MS run. It implements an abstract logic (right panel in **Fig.1**) which makes use of a decision tree, a common construct in computer science that has previously been applied in proteomics select optimal fragmentation modes by Coon and co-workers[35]. The decision tree simplifies the development of new acquisition strategies and generates a cascade of scan requests on the basis of the incoming scan containing the mass spectrum and the associated metadata. Every node of the scan protocol tree consists of three components: A filter, a scan template and data-dependent actions. The filter checks if meta- and spectral data of the incoming scan match particular features. If the check is negative, processing of this node and its children is stopped and the scan protocol tree proceeds to the next item. If the check is positive, for instance because the incoming scan is of MS1 type or contains specific ions of interest, then a new tailored scan request is created based on properties defined in the scan template. This comprises settings for the quadrupole, the collision cell and the mass analyzer. The third component, the data-dependent action, then establishes the connection between the incoming scan data and the settings of the next scan request. On the basis of its stored data and the incoming data, it chooses particular actions, such as selecting a particular precursor for isolation in the quadrupole, followed by acquisition of a fragmentation spectrum at a particular energy. Only the values that are different from the default template are overwritten. In the simple example of a topN method, the data-dependent action would be restricted to setting the position of the isolation window. After the incoming scan has successfully run through a scan protocol node, it is passed to its children, which may implement additional logic by themselves – such as checking for the presence of particular m/z values in the MS1 spectrum – and trigger further scan requests.

Although scan protocols allow an easy and flexible way to develop acquisition strategies on a high abstraction level, using them is complex and difficult for a non-specialized mass spectrometry laboratory. For this reason, MaxQuant.Live includes a series of small programs (apps) that can automatically generate scan protocols based on pre-defined acquisition strategies. We have created an app store for MaxQuant.Live that allows easy access to a collection of apps for different data acquisition strategies, which we have developed and tested in our group. In addition to the strategies described in this publication, BoxCar[36] acquisitions are already included and support for the EASI-tag method[37] is planned.

### Usability and performance of the software package

Our ambition was to make MaxQuant.Live very robust and fast, so that any mass spectrometry laboratory can use it for their workflows, without affecting ease of use or throughput. We further aimed to make it universally available and supportable in the long term, similar to the other parts of the MaxQuant ecosystem.

The graphical user interface (GUI) of MaxQuant.Live unifies the control over all our software components in a user-friendly way, starting from the instrument connection to the scan protocol library and the apps for creation of new acquisition schemes (**Fig. 2A**). The user can start a scan protocol from the library which then triggers MaxQuant.Live to take control of the mass spectrometer until the end of the run where it switches back to idle mode. It does not interfere with the vendor’s software and it can be continuously active in ‘listening mode’. In this way, acquisition can seamlessly switch between Xcalibur and MaxQuant.Live.

**Figure 2.**
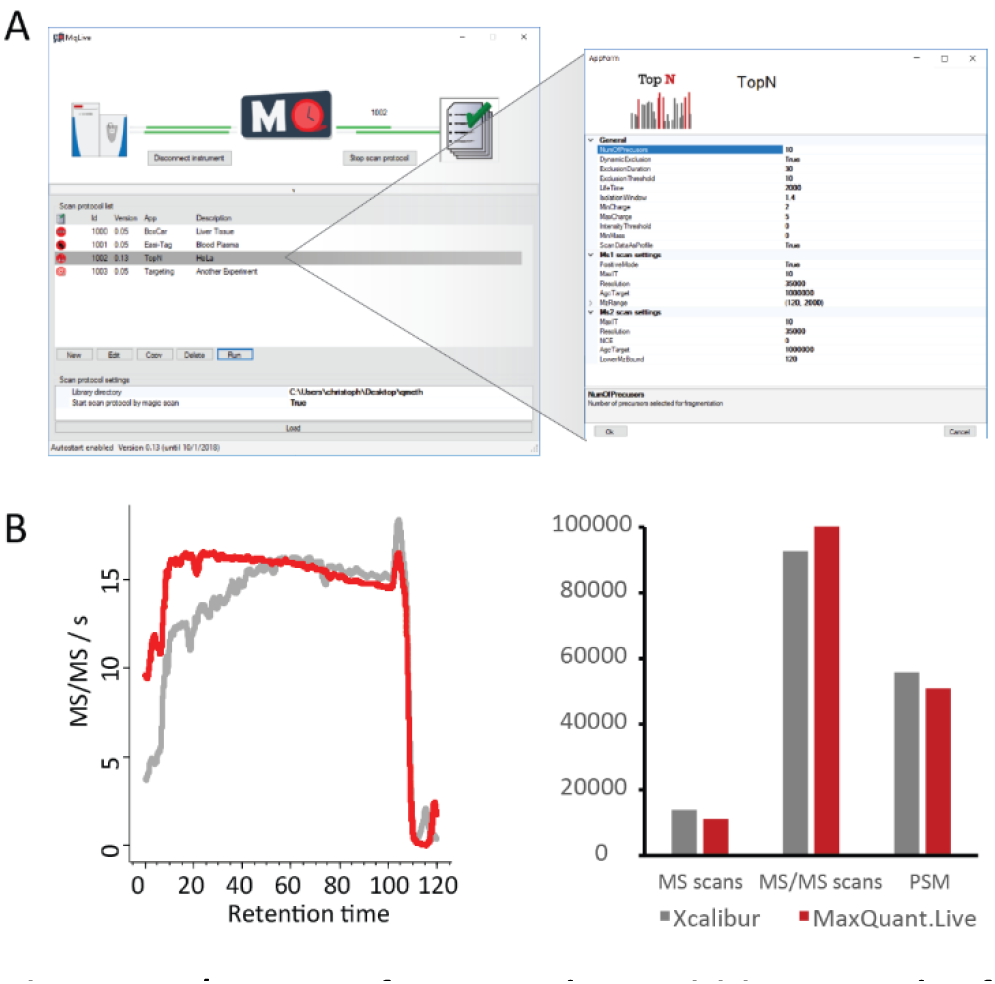
Ease of use and acquisition speed of MaxQuant.Live. A, Graphical user interface containing functionalities to create and manage scan protocols. B, Benchmarking the acquisition speeds of MaxQuant.Live vs. the vendor’s software (Xcalibur). In both top15 implementations, the instrument was acquiring MS2 spectra at nearly the maximum rate throughout the run (left panel). The number of MS and MS2 scans as well as the peptide spectrum matches (PSM) are comparable (right panel).

The user initially selects the app from the built-in app store for the desired workflow. MaxQuant.Live allows the creation of new scan protocols without knowledge of the underlying decision tree structure using simple GUIs that break down the complexity of each scan protocol into a small number of required settings. **Figure 2A** illustrates this for the GUI of the topN app, which re-implements the standard data-dependent acquisition scheme as a benchmark example. After the user specifies the parameters, the app generates the corresponding scan protocol and adds it to the scan protocol library. The GUI also allows editing a scan protocol within the app at later time points to modify the acquisition strategy.

In our topN implementation, the peak selection can be restricted to specific charge states and intensity values ranges to focus fragmentation on preferred classes. As in Xcalibur, re-sequencing of precursors can be prevented. Additionally, the relevant instrument parameters for the MS1 survey as well as the MS2 fragmentation scans can be specified in the GUI.

To benchmark the acquisition speed of a mass spectrometer under the control of MaxQuant.Live, we performed standard HeLa LC-MS/MS runs using our implementation of the top15 method. MaxQuant.Live achieved at least as many MS2 scans per second over the full 120 min gradient as the vendor’s software (**Fig. 2B**). (The faster speed at the beginning of the gradient is likely due to the fact that the Xcalibur peak selection algorithm uses a different intensity threshold.) This indicates that both MaxQuant.Live and the IAPI are extremely fast, and do not impose any relevant overhead in acquisition compared to direct control by Xcalibur. In particular, the total number of MS2 scans and peptide spectrum matches (PSMs) is not compromised, creating a solid basis for more intelligent acquisition schemes.

### Three dimensional adaptive control for peptide recognition in real-time

Having established a fast and robust framework for data-dependent acquisition, we next set out to accurately recognize eluting peptides in real-time at a very large scale. This is challenging because hundreds of precursor ions elute at any given time in complex proteome analysis and the elution time for every peptide subtly shifts from run to run. As a result, existing ‘inclusion lists’ and ‘exclusion lists’ are in practice limited to a relatively small number of precursors. In contrast, the ability to detect large numbers of peptides should enable MaxQuant.Live to take data-dependent decisions about the next scan operations in real-time and thereby to realize more intelligent acquisition strategies.

For this purpose, MaxQuant.Live includes a powerful app that implements diverse strategies for the targeting of specific precursor ions in an LC-MS run. They build on a real-time feature detection algorithm combined with adaptive non-linear corrections algorithm in the retention time, m/z scale and intensity dimensions.

In the list of centroid m/z values received from the instrument, our software determines isotope patterns, which are then compared to a list of precursor ions of know mass, charge state, intensity and estimated retention time. For a potential match, the ion intensity has to first exceed a threshold, which is a user-defined percentage of the expected intensity. This is calculated from the first two isotopic peaks, which are assumed to conform to the averagine-model[38], as is the case in MaxQuant post-processing, and which have to have the expected mass-to-charge difference within a user-defined tolerance of several ppm. MaxQuant.Live sets the recorded ion intensity to zero if either of the two peaks are missing or if there is an interfering peak before the presumed monoisotopic peak.

The second condition of the recognition algorithm requires that the precursor elutes within a certain time window around its expected retention time. Depending on a variety of external factors, peptide elution times can shift by several minutes between any two LC-MS runs, with the consequence that ‘retention time windows’ are generally set to several minutes. For very large numbers of targeted precursors, this would lead to too many potential matches to eluting features. To tackle this problem, we extended the recognition algorithm by an adaptive non-linear correction of the observed retention time shifts that is inspired by the ‘match between runs’ approach of MaxQuant[25]. Briefly, we use a subset of easily recognizable peptides to continuously minimize the median differences between the observed and the expected retention times. As a consequence of applying dynamic corrections to the ions properties we can dramatically shrink the tolerances for the elution time values that are used in the ion recognition. In a typical run, the interval containing 95% of the expected precursors shrinks from several minutes to less than one minute (**Fig. 3A**). While similar ‘dynamic corrections’ have been applied by us and others before, MaxQuant.Live achieves high robustness and precision by using a very large number of peptide precursors for real time correction (up to thousands, see below).

**Figure 3.**
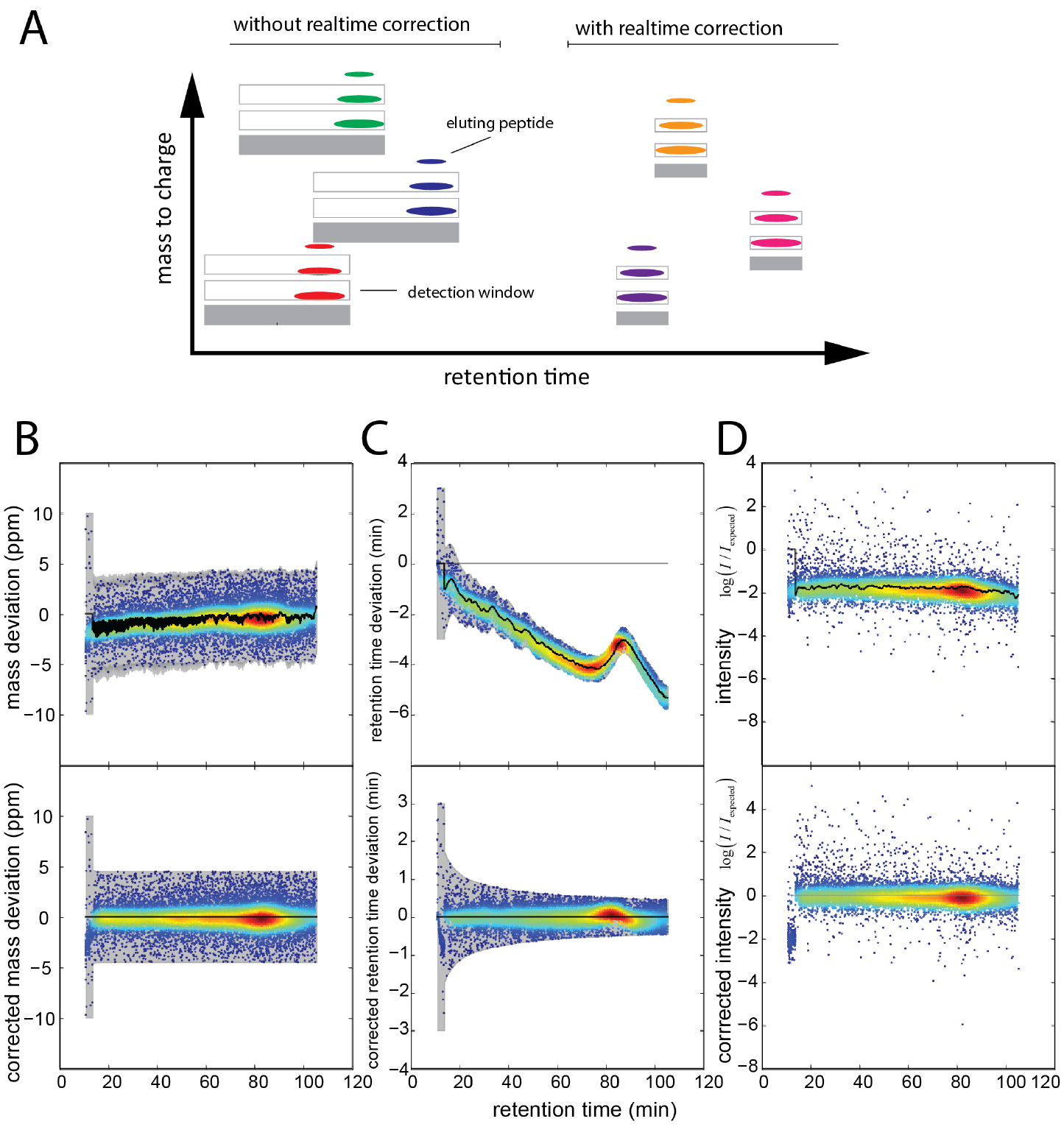
MaxQuant.Live targeting application. A, Real-time peptide recognition expects the first two isotopic peaks within a dynamic retention time and mass-to-charge tolerance window as indicated by the grey boxes. Our adaptive correction approach continuously corrects observed global shifts of the elution time, mass calibration and peptide intensity and reduces the tolerances to minimum values. B-D, application of the dynamic global corrections (black lines) during an LC-MS run (upper row) drastically narrows the recognition algorithm tolerances (gray areas in B and C) and the scaling of the peptide intensities to the values observed in the reference run.

Similar to the retention time alignment, mass accuracy can greatly be improved with the help of subsets of peptides that serve as internal calibrants[39]. Based on the same principle as above, we therefore continuously recalibrate the mass scale, achieving several fold improvements in real time mass accuracy. However, in contrast to MaxQuant, our mass correction applies to each entire spectrum, rather than being peptide specific. For the example in **Fig. 3B**, the a priory mass window could be reduced from a maximum mass deviation ±10 ppm to ±4.5 ppm which is the same maximum value as used in MaxQuant post-processing.

Signal intensity is the third dimension of precursor features and its adaptive control accounts for day to day differences in sensitivity of the LC-MS set up. Given the signals of the reference peptide population, an overall scaling factor is applied to make the recorded signal intensities comparable to the ones in the targeting list generated from a reference run. In our experiments we noticed that this scaling factor varied between different runs, for example as a result of varying sample amounts on column,, but only little within a single run (**Fig. 3C**).

### Targeted acquisition of breakdown curves

Robust and precise peptide recognition in real-time should open up various opportunities for advanced analysis of selected peptides. To demonstrate this, we chose to generate ‘breakdown curves’, which are useful to determine optimal collision energies of peptides or to determine the structure of metabolites. We directed MaxQuant.Live to detect a subset of 1,000 peptides in a complex HeLa background and fragment each of them with increasing collision energies. Using 10,000 abundant background peptides for our adaptive real-time correction, the monitoring time for each of the peptides of interest was reduced to less than 4 min in the 120 min runs. Notably, the median absolute retention time deviation from the original targeting list was up to 4 to 6 min before, and only 0.6 min after re-calibration in all three replicates (***Suppl. Fig. 1***). Together with the sub-ppm mass accuracy, this allowed us to successfully acquire breakdown curves for 818 of the 1000 targeted peptides. **Figure 4a** illustrates the method for a specific target peptide (EQQIVIQSSGGLSK). MaxQuant.Live detected the elution peak about 5min before it was expected given the uncorrected retention time and -upon surpassing the normalized intensity threshold-scheduled the peptide for several breakdown analyses (one example shown in **Fig. 4a**). We utilized ten different collision energies from NCE 18 to 36 at a mass resolution of 15,000 at *m/z* 200, which translates into a net analysis time of only 0.3 s per breakdown curve. Three example spectra for low, middle and high collision energies are annotated in **Fig. 4b**. At NCE 18, the spectrum was dominated by the precursor ion, indicating incomplete fragmentation. Despite the relatively low abundance of fragment ions, we observed an almost complete y ion series (*y_1_* – *y_11_*) as well as the complementary *b_3_* to *b_7_* ion series. For the *b* ion series, we observed prominent neutral loss peaks at an NCE to 26. Increasing the NCE further yielded many low-mass immonium ions and internal fragment ions, which escaped automatic scoring with the Andromeda search engine[40]. The possibility to target thousands of peptides enables global analysis of peptide fragmentation. To illustrate uses of this capability, we plotted the fraction of the fragment ion current that has been identified as a function of the collision energy (**Fig. 4c**). This value peaked at NCE 22-24, presumably due to the less frequent generation of internal fragment ions. Generally, we noted a wide distribution for the peptide specific optimal NCEs, highlighting sequence-dependent differences in the fragmentation efficiency even with the normalized collision energies. Next, we investigated the energy-dependent generation of b and y ions (**Fig. 4d**). B ions were preferably generated at lower collision energies, while the number of annotated y ions (including neutral losses) increased with higher collision energy. About 50% of the ion current was accounted for by y ions throughout all NCEs, while the relative abundance of b ions was decreasing (**Fig. 4e**). Interestingly, the fraction of the a2 fragment ion in the characteristic a2, b2 ion pair that is formed instead of the b1 ion, increased up to a full 20% of the annotated fragment ion current at higher collision energies.

**Figure 4.**
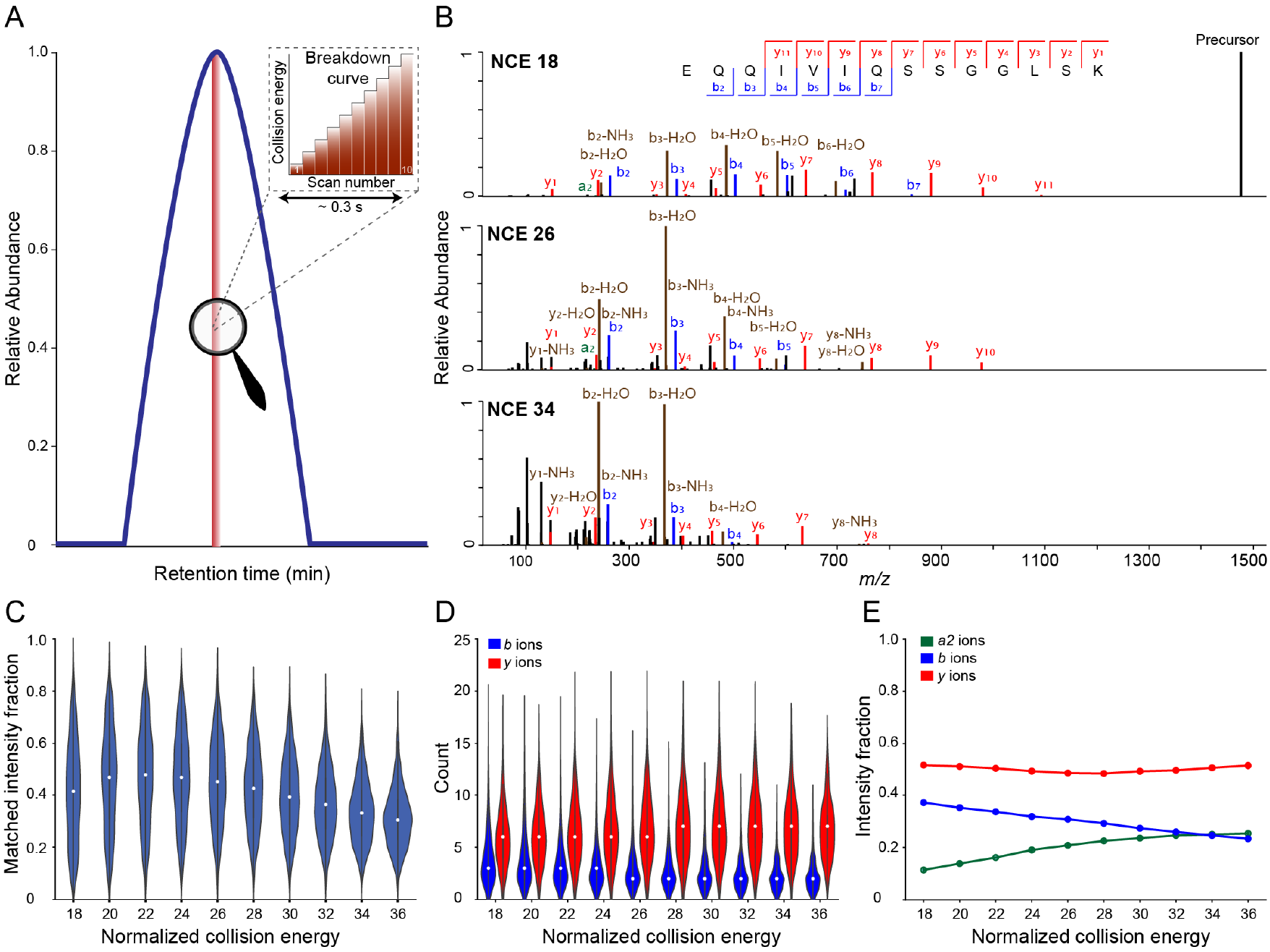
Automated acquisition of peptide breakdown curves. A, Extracted ion chromatogram of a targeted peptide. Upon detection, MaxQuant.Live acquires repeated MS2 scans of this precursor with increasing collision energies. B, Exemplary spectra from a single breakdown curve. C, Fraction of the total MS2 ion current annotated by the Andromeda search engine as a function of the normalized collision energy. D, Number of identified b and y ions as a function of the normalized collision energy (NCE). E, Summed intensity of a2, b, and y ions relative to the sum of all identified fragment ions. N = 818 peptides.

### Predictive Multiplexed Selective Ion Monitoring (pmSIM)

In the example above, the MS1 signal of a targeted peptide triggered the acquisition of MS2 scans. However, MS1 spectra can be incomplete in that low abundance precursors may be present in some but not other runs. Thus, instead of relying on the MS1 trigger signal and motivated by the high accuracy of the real-time retention time alignment described above, we predicted the elution of target peptides based on the endogenous background population (**Fig. 5A**).

**Figure 5.**
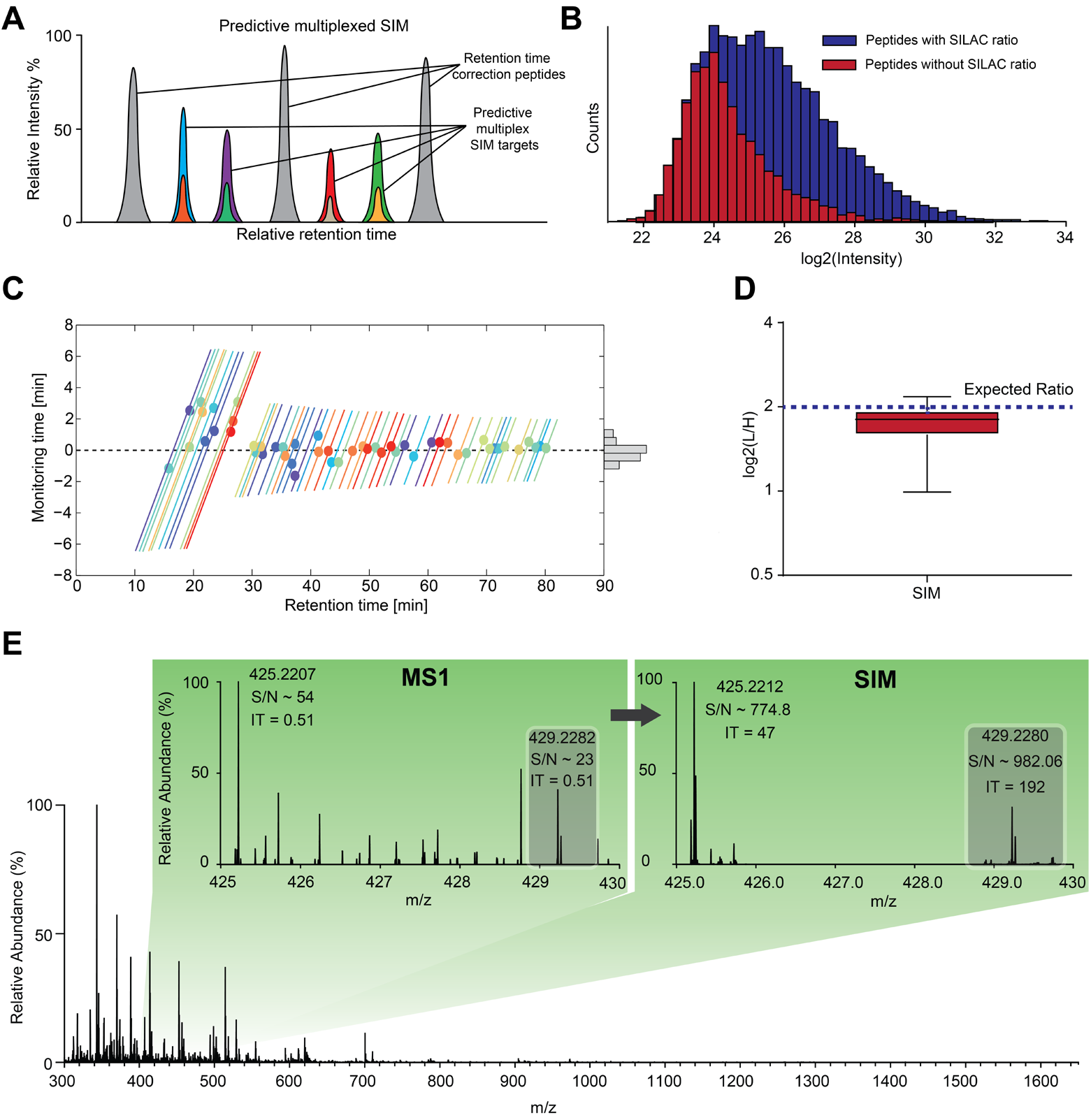
Predictive multiplexed SIM (pmSIM). Peptides employed for live retention time correction are distributed across the gradient and are highly abundant, compared to the low abundant targeted peptides and their SILAC partner needed for quantification (A). B: Total log2 intensity abundance range of all peptide identifications from the standard run with a light to heavy SILAC ratio of 4:1. In blue, all peptide identifications with identified ratios are highlighted. In red, all peptide identifications in the light channel without any reported SILAC ratios are highlighted. C: Watch-time of the predictive multiplexed SIM scan-mode in the targeting experiment. D: Resulting SILAC ratios after SIM-targeting using Skyline [32]. E: Example of a very low abundant target peptide compared in the original MS1 and after pmSIM demonstrating ^~^40-fold increase of S/N.

To demonstrate our approach, we set up a SILAC[31] experiment in which heavy and light whole-cell HeLa digests were mixed in a 1:4 ratio. In DDA, the limited dynamic range of the full scan resulted in many missing heavy-to-light ratios in the low intensity range (**Fig. 5B**) and overall, MaxQuant reported ratios for only ^~^60% of the identified peptides. The sensitivity can be boosted dramatically by isolating and selectively accumulating narrow *m/z* ranges, which results in improved MS1 (SIM) or, when fragmented, MS2 quantification (termed parallel reaction monitoring ‘PRM’, when used in targeting studies[13]). Without adaptive retention time alignment, such scans have to be repeated over a time range large enough to account for the typical shifts and fluctuations in the elution times. This typically results in an very large over-head of scans for each targeted peptide, limiting the total number that can be studied in a single LC-MS run. Here, we used a “predictive multiplexed SIM” (pmSIM) method to measure heavy and light SILAC peptides simultaneously.

We again selected 10,000 high abundant peptides as correction peptides for the adaptive real-time corrections (**Fig. 5A**). The correction algorithm of MaxQuant.Live dynamically centered the observation time ranges around the peptide elution times (**Fig. 5C**, colored lines and circles, respectively), yielded an accurate prediction of the time range in which each peptide was expected to elute. This resulted in three times smaller window sizes compared to the initial values. The comparison of the window sizes with the deviations of the peptide apex times from the predictions (**Fig. 5C**, histogram) shows that the time windows could even have been chosen smaller. It should be noted that our settings were very conservative and the number of target peptides could be much higher than the factor three already demonstrated here.

To validate the accuracy of our prediction algorithm, we targeted 50 peptides from the low abundance range with missing ratios from our SILAC HeLa study (**Fig. 5B**). We then used the MaxQuant.Live targeting app to specify an acquisition method that executed SIM scans of the corresponding ion pairs repeatedly over the expected elution time range. The pmSIM strategy correctly quantified the ratios for the targeted peptides close to the expected value of 1:4 (**Fig. 5D**). This is notable, because none of the heavy labelled peptides was detectable at the MS1 level before. The example in **Fig. 5E** shows the increase in sensitivity by comparing the MS1 with the corresponding SIM scan. In the SIM scan, the injection time for the previously unrecorded heavy peptides is 400 times larger than the injection time of the full scan, drastically improving the quantitative accuracy.

### Global targeting for efficient proteome quantification

The examples shown so far demonstrate the ability of MaxQuant.Live to perform a specific and sophisticated analysis of a limited number of peptides of interest. The fact that the underlying peptide recognition algorithm can in principle deal with an unlimited number of peptides, makes applications feasible that target a substantial proportion of the total set of precursor ions. We term this approach ‘global targeting’ and reasoned that it should generically boost the reproducibility of peptide fragmentation events between LC-MS runs compared to the topN method.

We implemented global targeting using the targeting app of MaxQuant.Live to investigate the potential benefits of this approach. To do this, we generated sets with different numbers of targeted peptides, which were randomly selected from triplicate MS analysis of tryptic HeLa lysates using a standard top15 method in Xcalibur (**Fig. 6A**).

For every set of peptides we performed triplicate global targeting LC-MS runs in which MS2 scans were triggered if one of the peptides was recognized by our algorithm in the MS1 scans. The number of peptides that were fragmented and correctly identified afterwards by MaxQuant is shown in **Figure 6B** for all six sets of peptides. While nearly every targeted peptide was hit in at least one of the runs (green line), this fraction decreases for those hit in at least two (blue line) and all three runs, respectively (red line). This indicates that our strategy is capable of targeting peptide numbers which are orders of magnitude higher than the current state of the art in routine practice, even though some stochasticity remains between the acquisitions in the different runs. This is likely due to some peptides not being recognized by our algorithm at the MS1 level. Surprisingly, an analysis of the of the initial topN and the global targeting raw data files using MaxQuant with the matching between runs feature showed very similar results. Thus, the well-established feature detection of MaxQuant could not find significantly more spectral features at the MS1 level from the targeting list, even given full information after complete analysis. This suggests that these peptides, which were selected from the original topN runs, are not “visible” at all in the MS1 scans of the targeting runs, which were measured days later. A comparison of the intensity histograms of the peptides that were successfully targeted and identified by MS/MS in all three runs (**Fig. 6C**, blue) to the ones that were only hit in one (green) shows a slight shift of the intensity distribution to lower values. We therfore suspect that the spectral noise thresholding employed by Xcalibur set the corresponding signals to zero even though they only slightly dipped below acceptance criteria, something we have also noticed when boosting the dynamic range in the BoxCar acquisition scheme[36]. A possible solution to tackle this problem of the acquisition side would be to boost the peptide intensities by using BoxCar[36] scans for the peptide recognition instead of the MS1 scans.

**Figure 6.**
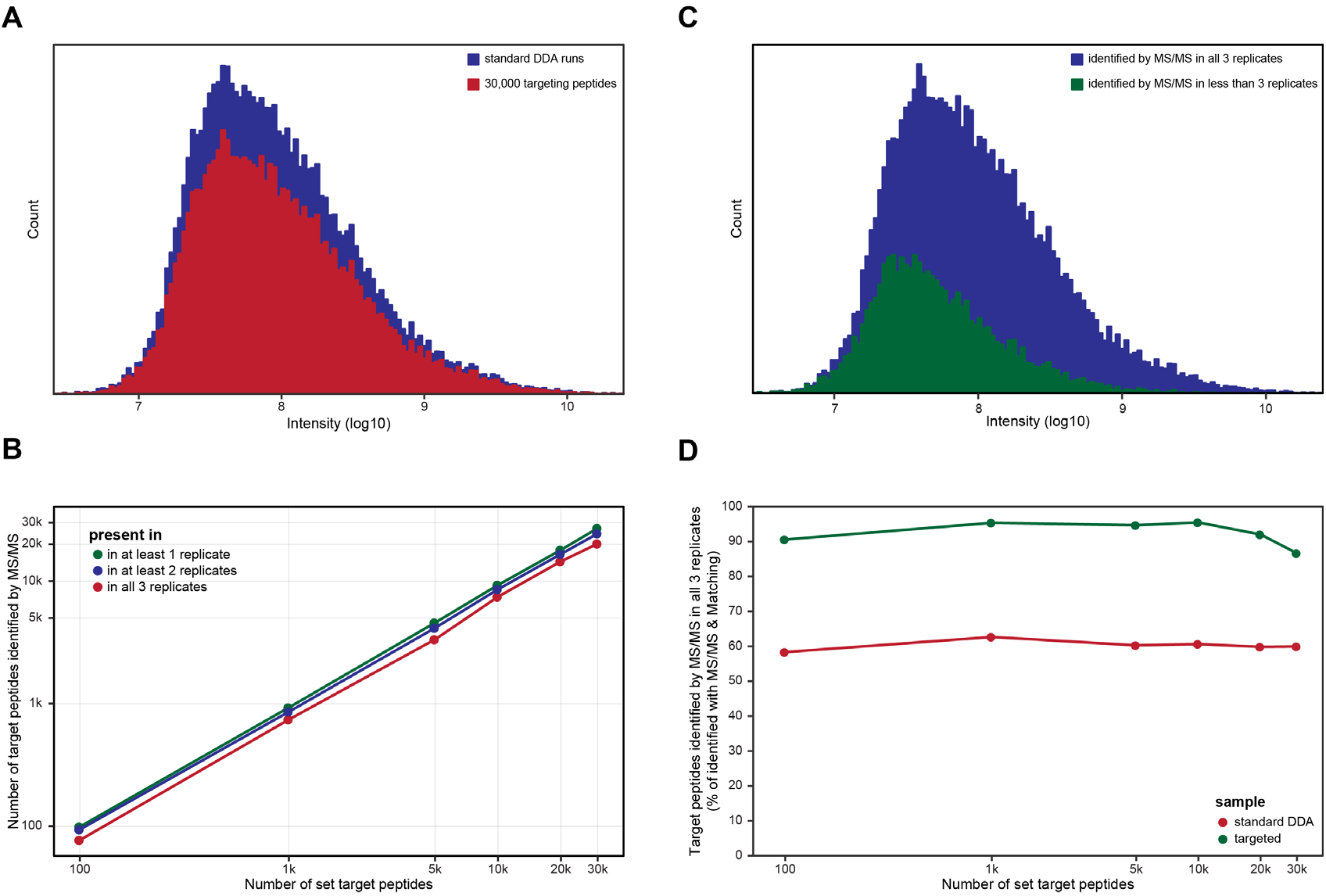
Global targeting allows reproducible identification of at least 20,000 peptides. Tryptic HeLa lysates were analyzed by either a standard DDA method designed in the Xcalibur method editor or by the MaxQuant.Live targeting method with 100, 1,000, 5,000, 10,000, 20,000 or 30,000 targeted peptides. Triplicate injections were performed and sequenced peptides were identified by MaxQuant with or without the matching between runs function activated. (A) Selected targets were uniformly distributed over the whole intensity range of peptides identified in previous triplicate standard DDA runs. (B) The number of targets correctly identified by MS/MS. (C) Intensity distribution of the correctly identified sequences in MaxQuant.Live targeting runs with 30,000 targets. (D) Percentage of correctly identified targets by MS/MS for the MaxQuant.Live targeting method compared to the identifications by MS/MS in the standard topN method and matching between runs in all three triplicate injections.

To counter the effect of peptide features missing due to thresholding in our analysis and make the data between the two triplicate measurements comparable, we further normalized the number of peptides that were successfully targeted in all triplicate measurements to the number of targeted peptides identified by the matching between run feature. The normalized percentage of hit target peptides in the targeting runs (green line) shows a success rate of about 90% for nearly all the targeting sets but drops down to 85% for the ones containing more than 20,000 peptides. This effect is related to the fact that the number of co-eluting peptides increases beyond what can be fragmented sequentially by the mass spectrometer in the time available. Some of the peptides are then fragmented too late after their elution maximum and the respective spectra MS2 spectra are therefore of too low intensity. These are not limitations of the global targeting approach, but opportunities to be addressed by instrument improvements. In any case, the advantages of global targeting become obvious if one compares these results with the normalized percentage of the targeted peptides fragmented in all topN triplicates (red line). Instead of the 90% success rate of global targeting, the standard method only reached about 60%, regardless of the number of peptide precursors considered here.

## DISCUSSION AND CONCLUSION

In bottom up proteomics, data dependent acquisition and targeted approaches have co-existed for many years. DDA has been and remains the method of choice for initial characterization of proteomes under study. Conversely, there are many applications, where only a restricted number of peptides is of interest, but these need to be measured consistently over many samples. Although both approaches have become more powerful with the general advances in instrumentation and proteomics technologies, DDA is still not powerful enough to subsume targeted analyses. Conversely, targeted method have been difficult to establish in a robust manner, sufficient for clinical use, for instance, especially when monitoring more than a few dozens of peptides. As a result, increasingly powerful DIA methods have become alternatives to targeted proteomics in some applilcations[11]. Beyond these general considerations there is also a large unexplored territory of sophisticated analysis of a subset of peptides, which could not be addressed by any of these methods so far.

Here, we made use of the recently developed fast and robust IAPI of the Thermo instruments to interface with the acquisition process in real time. Our overall goal was to make use of experimental information as they are acquired, in order to direct the acquisition in a more intelligent way. We achieved this by writing an application that implements different scan modes in the form of small built-in ‘apps’, each for a class of mass spectrometric acquisition schemes. We demonstrated that this workflow is highly performant as it can easily replicate the standard topN methods, for instance, without loss of quality. MaxQuant.Live continuously recalibrates the signal coming from the mass spectrometer in retention time, mass and intensity dimensions, allowing a much better prediction of the identity of eluting peptide features than possible previously. This allows selecting any subgroup of hundreds of peptides to be targeted for accurate quantification (exemplified by our predictive SIM targeting) or special acquisition methods. In depth analysis of the fragmentation patterns of large numbers of individual peptides is a valuable addition to the proteomics toolbox, for instance to optimize precursor-fragment transitions in targeted analyses or to pin-point and localize modifications of low-abundance proteins. MaxQuant.Live ensures that all peptides are reliably acquired at all collision energies (as opposed to stochastic precursor selection with DDA) and in single runs. Even if these methods take much longer than standard fragmentation for the selected peptides, they still do not substantially contribute to overall measuring time. This means that sophisticated measurements could be done on peptides of interest, while still recording the overall proteome. While we randomly chose peptides, one could, for instance, select a specific class of post-translation modifications, peptides that distinguish between isoforms or any other highly informative class of interest. The resulting, ‘enhanced’, datasets could also become valuable sources for imminent machine-learning approaches[18], [26]. In our own group, we have already applied such a strategy to characterize the fragmentation of a novel isobaric tag[37].

Finally, building on the precise re-calibration in MaxQuant.Live, we demonstrate that the scale of such experiments can be readily extended to over 20,000 target peptides with high reproducibility. The global targeting approach thus allows to acquire previously unavailable data even on large subsets of the proteome under study. We have made the tools freely available and hope that they will stimulate the community into exploring this exciting direction.

## ACKNOWLEDGEMENTS

We thank our colleagues in the Max-Planck Institute of Biochemistry and at Thermo Fisher Scientific for discussion and help, in particular Drs. A. Kreutzmann, D. Mourad, K. Ayzikov, F. Grosse-Coosmann, S. Horning and A. Makarov. This project has received funding from the European Union′s Horizon 2020 research and innovation programme under grant agreement no. 686547, by the German Research Foundation (DFG–Gottfried Wilhelm Leibniz Prize) granted to Matthias Mann and by the Max-Planck Society for the Advancement of Science.

